# V367F mutation in SARS-CoV-2 spike RBD emerging during the early transmission phase enhances viral infectivity through increased human ACE2 receptor binding affinity

**DOI:** 10.1101/2020.03.15.991844

**Authors:** Junxian Ou, Zhonghua Zhou, Ruixue Dai, Shan Zhao, Xiaowei Wu, Jing Zhang, Wendong Lan, Lilian Cui, Jianguo Wu, Donald Seto, James Chodosh, Gong Zhang, Qiwei Zhang

## Abstract

The current global pandemic of COVID-19 is caused by a novel coronavirus SARS-CoV-2. The SARS-CoV-2 spike protein receptor-binding domain (RBD) is the critical determinant of viral tropism and infectivity. To investigate whether naturally occurring mutations in the RBD during the early transmission phase have altered the receptor binding affinity and infectivity, firstly we analyzed *in silico* the binding dynamics between mutated SARS-CoV-2 RBDs and the human ACE2 receptor. Among 32,123 genomes of SARS-CoV-2 isolates (January through March, 2020), 302 non-synonymous RBD mutants were identified and clustered into 96 mutant types. The six dominant mutations were analyzed applying molecular dynamics simulations. The mutant type V367F continuously circulating worldwide displayed higher binding affinity to human ACE2 due to the enhanced structural stabilization of the RBD beta-sheet scaffold. The increased infectivity of V367 mutants was further validated by performing receptor-ligand binding ELISA, surface plasmon resonance, and pseudotyped virus assays. Genome phylogenetic analysis of V367F mutants showed that during the early transmission phase, most V367F mutants clustered more closely with the SARS-CoV-2 prototype strain than the dual-mutation variants (V367F + D614G) which emerged later and formed a distinct sub-cluster. The analysis of critical RBD mutations provides further insights into the evolutionary trajectory of SARS-CoV-2 under negative selection pressure and supports the continuing surveillance of spike mutations to aid in the development of new COVID-19 drugs and vaccines.

**Importance:** A novel coronavirus SARS-CoV-2 has caused the pandemic of COVID-19. The origin of SARS-CoV-2 was associated with zoonotic infections. The spike protein receptor-binding domain (RBD) is identified as the critical determinant of viral tropism and infectivity. Thus, whether the mutations in the RBD of the circulating SARS-CoV-2 isolates have altered the receptor binding affinity and caused them more infectious, should be paid more attentions to. Given that SARS-CoV-2 is a novel coronavirus, the significance of our research is in identifying and validating the RBD mutant types emerging during the early transmission phase that have increased human ACE2 receptor binding affinity and infectivity. The RBD mutation analysis provides insights into SARS-CoV-2 evolution. The continuous surveillance of RBD mutations with increased human ACE2 affinity in human or other animals is important and necessary, particularly when the direct correlation between the virus variations and vaccine effectiveness is underdetermined during the sustained COVID-19 pandemic.

## 1. Introduction

A novel coronavirus SARS-CoV-2 has caused outbreaks of Coronavirus Disease 2019 (COVID-19) globally beginning in mid-December 2019, with an epicenter in Wuhan, China(1–3) As of November 22, 2020, SARS-CoV-2 had infected 57.8 million people world-wide and caused 1.3 million deaths with an estimated fatality rate of 2.25%(4). This on-going pandemic of COVID-19 has become the most serious threat to public health in this century.

The origin of SARS-CoV-2 remains elusive. However, the initial cases were largely associated with a seafood market, which indicated potential zoonotic transmissions(1, 5, 6). Although bats and pangolins are most likely the reservoir and the intermediate hosts in the wild, more evidence is needed to support zoonotic transmission and to track the origin of this new coronavirus(5, 6).

Angiotensin-converting enzyme 2 (ACE2) is the host cellular receptor for the SARS-CoV-2 spike glycoprotein, which is similar to its counterpart in SARS-CoV. The receptor-binding domain (RBD) of the spike protein subunit S1 interacts directly with ACE2, providing for tight binding to the peptidase domain of ACE2(7–10). Therefore, RBD is the critical determinant of virus-receptor interaction and reflects viral host range, tropism, and infectivity. Although the RBD sequences of different SARS-CoV-2 strains circulating globally are largely conserved, mutations have appeared; these may account for differences in viral infectivity and also contribute to its spread(10–14).

Meanwhile, S glycoprotein participates in antigenic recognition and is expressed on its protein surface, likely to be immunogenic as for carrying both T-cell and B-cell epitopes. The potential antibody binding sites that have been identified indicates RBD has important B-cell epitopes. The main antibody binding sites substantially overlap with RBD, and the antibody binding to these sites is able to block viral entry into cells(14–16).

To investigate whether mutations in RBD emergent during the early transmission phase have altered the receptor binding affinities and whether these isolates may have been selected for higher infectivity, the binding dynamics and the infectivity between the SARS-CoV-2 RBD mutants and human ACE2 receptor were modeled and assessed computationally. The evolutionary trajectory of SARS-CoV-2 during the early transmission phase under negative selection pressure was also analyzed. In addition, experimental validation of the enhanced affinity and infectivity of the V367F mutant was performed.

## 2. Results and discussion

### 2.1 SARS-CoV-2 Spike and RBD mutation mapping and scanning during the early transmission phase

32,123 SARS-CoV-2 isolates with whole-genome sequences available in public databases and sampled before March 31, 2020 were analyzed. Among them, 302 isolates with RBD mutations were identified when compared with reference strain Wuhun-Hu-1 that was the firstly isolated in Wuhan in December 2019. All the mutants were clustered into 96 mutant types, six of which were dominant mutant types that were found in more than ten isolates (Table 1): V483A (35x), V367F (34x), V341I (23x), N439K (16x), A344S (15x), G476S (12x) (Figure 2C). V483A and V367F accounted for 11.59% and 11.26% of 302 mutants, respectively. V367F mutants occurred on Jan 22, 2020, which was the earliest dominant mutant type. During the early transmission phase, there was no D614G mutation in 35 V483A mutants; however, there were 11 of 34 V367F mutants that harbored D614G mutation. The detailed alignment of amino acid sequences in the RBD of the mutants were shown in Supplementary Figure 1. These mutants were emergent in multiple continents, including Asia, Europe, North America and Oceania. Most RBD mutants were circulating in Europe and North America (Figure. 1). V367F mutants were found in all the four continents.

**Table 1.** Dominant mutations in the spike RBD (>10 isolates) and their binding affinity change (as of March 31, 2020). The numbers of dominant mutations in the spike RBD (as of March 31, 2020 collected) are shown. The date of mutant occurrence, ID of the first mutant, and concurrence of D614 or G614 mutants are identified. V483A mutants (35x) were only found in D614 isolates. V367F mutants were first identified in D614 isolates (23x), then identified in G614 mutants (11x). Receptor binding efficiencies of dominant mutants were evaluated by both amino acid property and binding free energy change using the MM-PBSA method (18, 19). The number of total mutants are 302.

| Amino acid<br>position in<br>spike gene | Amino<br>acid<br>change | Percentage<br>of mutants | Date of<br>occurrence | ID of the first<br>mutant | D614 | G614 | Total | Binding free<br>energy<br>increase |
| --- | --- | --- | --- | --- | --- | --- | --- | --- |
| 483 | V to A | 11.59% | 2020/2/29 | EPI_ISL_417159 | 35 | 0 | 35 | No |
| 367 | V to F | 11.26% | 2020/1/22 | EPI_ISL_408975 | 23 | 11 | 34 | Yes |
| 341 | V to I | 7.95% | 2020/3/3 | EPI_ISL_454450 | 0 | 23 | 23 | No |
| 439 | N to K | 5.30% | 2020/3/16 | EPI_ISL_425684 | 0 | 16 | 16 | No |
| 344 | A to S | 4.97% | 2020/3/7 | EPI_ISL_506954 | 2 | 13 | 15 | No |
| 476 | G to S | 3.97% | 2020/3/2 | EPI_ISL_417081 | 2 | 10 | 12 | No |

**Fig. 1:**
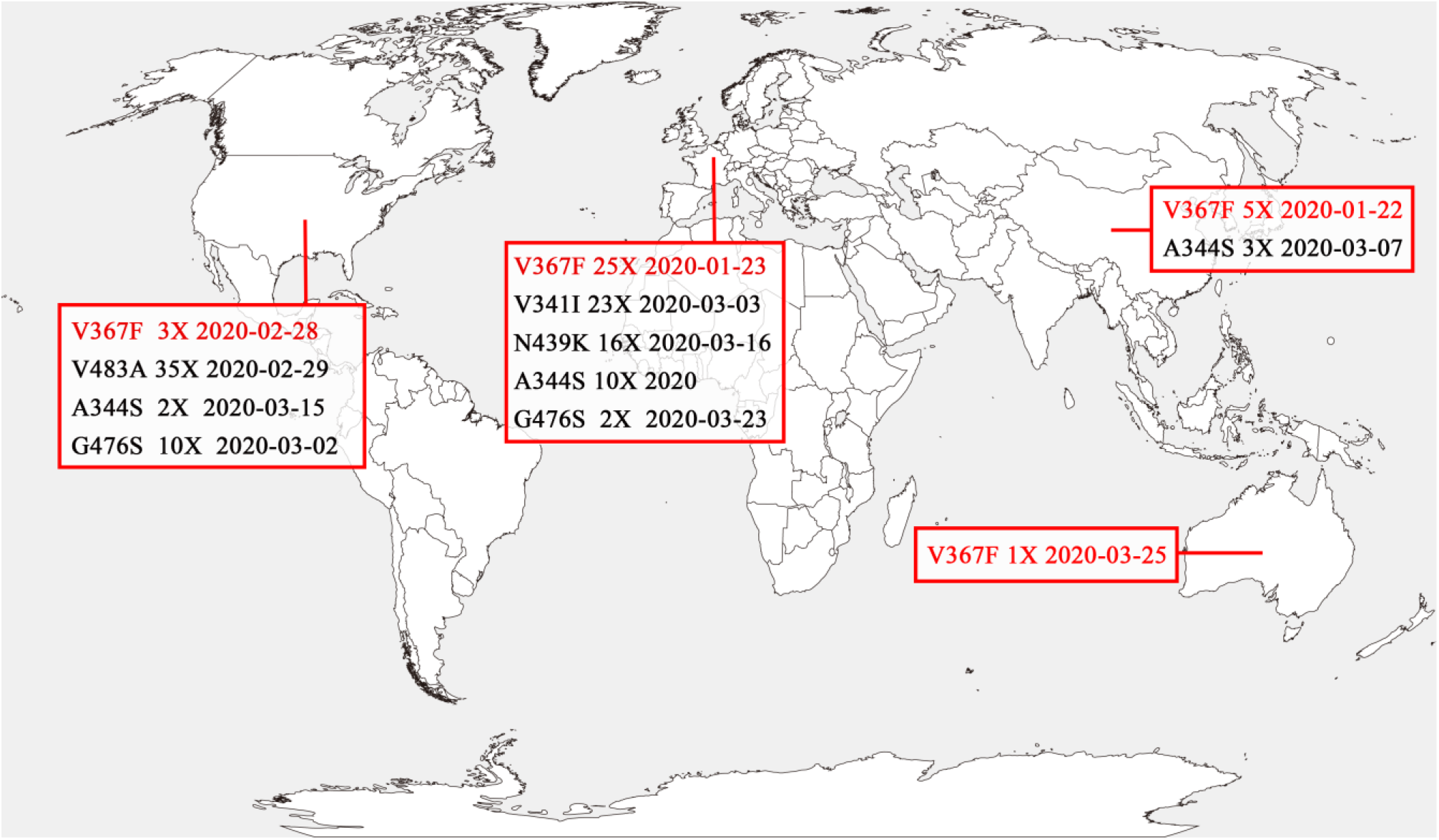
Geographical Distribution of the SARS-CoV-2 RBD mutants. The geographic distribution of the RBD mutants in four continents (>10 isolates) is displayed. The mutants marked in black are mutants with similar binding affinities with strain Wuhan-Hu-1. V367F mutants with the enhanced binding affinity were found in all the four continents and marked in red. The mutants analyzed were isolated as of March 31, 2020.

**Fig. 2.**
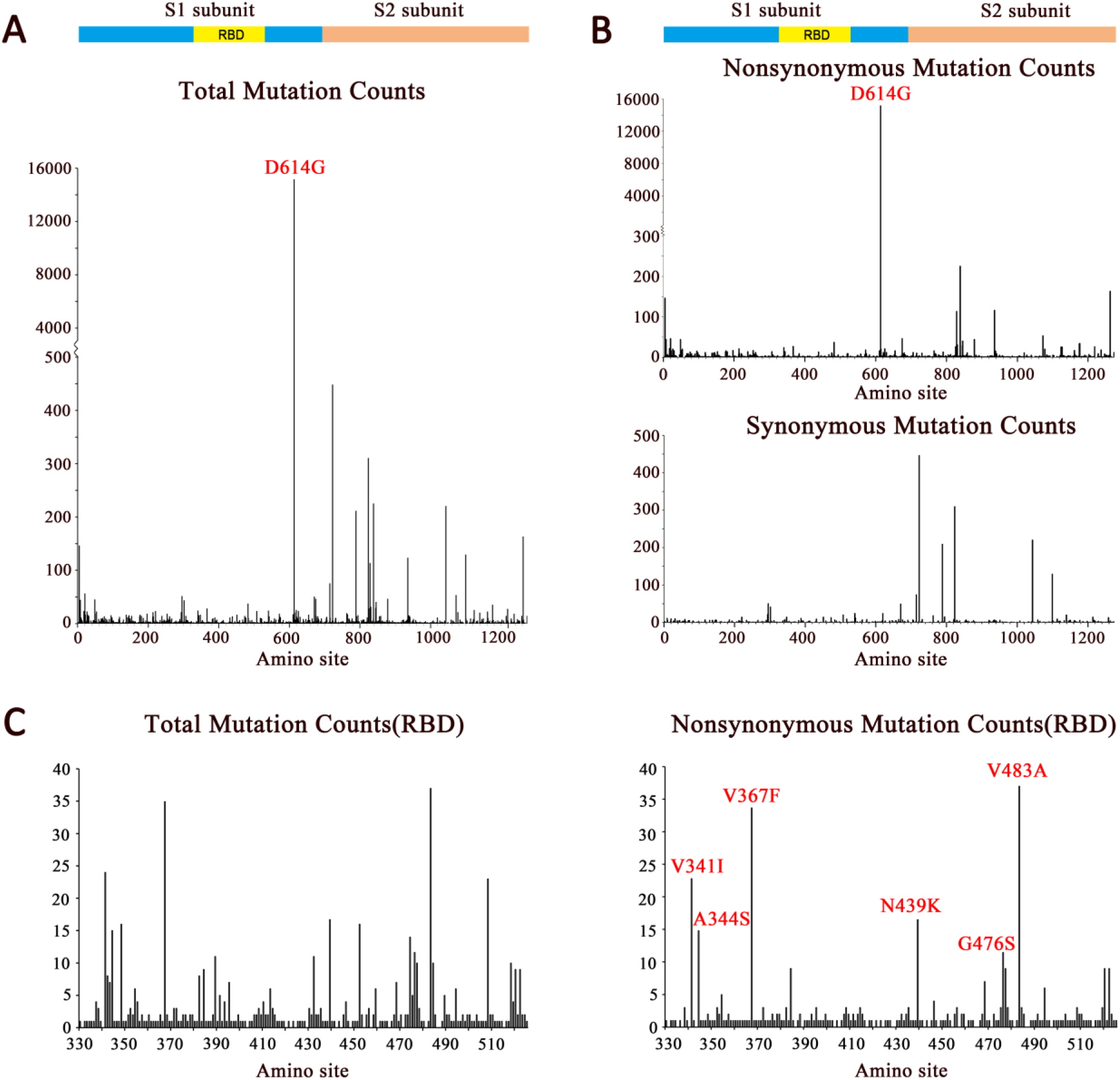
Mutation scanning graph of SARS-CoV-2 S gene. Mutation sites comparing to the strain Wuhun-Hu-1 were analyzed by BioAider (V1.314) (17). **(A)** The total mutation (synonymous and nonsynonymous) in spike gene; **(B)** The nonsynonymous and synonymous mutations in spike gene; **(C)** The total mutations and nonsynonymous mutations in RBD. All positions containing gaps and missing data were eliminated. Structural domains are annotated. The peak signal of D614G and the other dominant mutations are marked in red. The ordinate showed the number of the mutants. The isolates analyzed were isolated as of March 31, 2020.

### 2.2 Nucleotide diversity and selective pressure on SARS-CoV-2 spike gene and RBD

As SARS-CoV-2 S protein mediates attachment of the virus to cell-surface receptors and fusion between virus and cell membranes, the polymorphism and divergence of S gene were analyzed by DnaSP6 (version 6.12.03) (15). Overall, S2 subunit was more diverse than S1 subunit (Figure 2A). D614G mutation in S1 subunit accounted for most of the polymorphism and divergence. Most of the other synonymous and nonsynonymous mutations occurred in S2 subunit (Figure 2B), of which synonymous mutations were more than nonsynonymous mutations. During the early transmission phase, RBD was conserved and not as diverse as S2 subunit (Figure 2A and 2B). The six major mutant types were shown in Figure 2C.

Since RBD is the only domain to bind human ACE2 receptor and, in turn, initiates cell entry, it is believed that the RBD should be highly conserved. To test this hypothesis, we investigated the selective pressures of the S gene during the early transmission phrase by calculating nonsynonymous/synonymous substitution rate ratios (dN/dS ratios) for various segments of the S gene from the 32,123 SARS-CoV-2 isolates (20, 21). The entire S gene exhibited a dN/dS of 0.86, close to 0.83 reported previously (22), showing that the S gene is indeed under negative selection (Table 2). However, different regions of the S gene may be subject to different selection pressures. The S1 gene exhibited unusually high dN/dS (2.05) due to the widely spreading non-synonymous mutation D614G in the S1 gene (23). The S2 gene exhibited a lower dN/dS (0.2388551), indicating that the S2 gene was more conserved than S1 under negative selection. The S1-RBD showed lower dN/dS (0.7258567) compared to the entire S gene (0.8569338). Therefore, the functional relevance of the RBD mutations may be inferred.

**Table 2:**
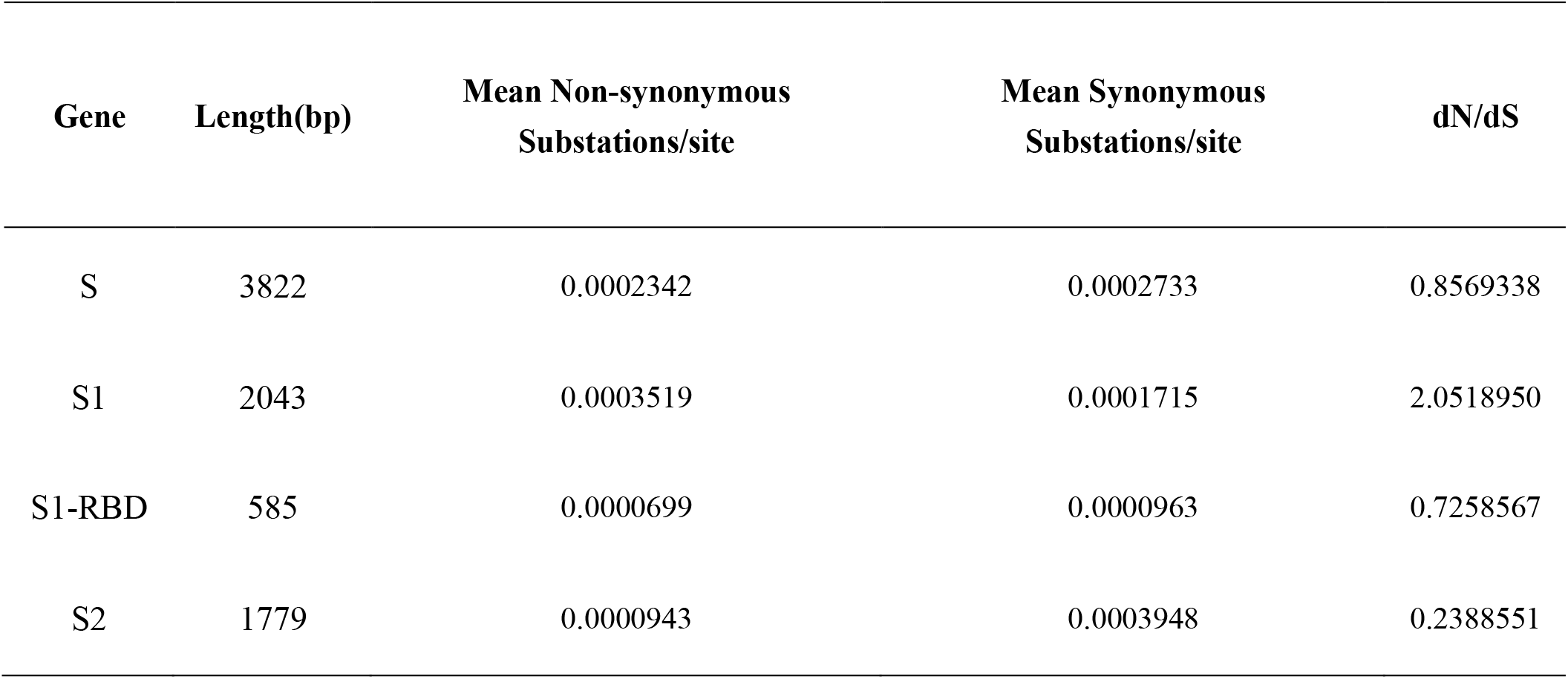
Nucleotide substitution rates and selection pressures for S gene. The numbers of nonsynonymous and synonymous differences per sequence from averaging over all sequence pairs are shown. Analyses were conducted using the Nei-Gojobori method(Jukes-cantor model) in Mega X (10.0.2) (20, 21). The analyses involved 32,123 SARS-CoV-2 S gene sequences. All positions containing gaps and missing data were discarded.

### 2.3 V367F mutant emergent during the early transmission phase binds human ACE2 receptor with higher affinity

To estimate the functional changes suggested by the RBD mutations, we performed molecular dynamics (MD) simulations for the prototype SARS-CoV-2 (Wuhan-Hu-1 strain) and the RBD mutants in order to assess their binding energy to human ACE2 receptor. Each simulation was performed at 10ns and each model was simulated in triplicate. All trajectories reached a plateau of RMSD after 2~5ns (Figure 3A), indicating that their structures reached equilibrium. All of the subsequent computations on thermodynamics were based on the 5~10ns trajectories. The ΔG of V367F mutant was significantly low (~ 60 kJ/mol) (*P*=0.0151), approximately 25% lower than the prototype strain (−46.5 kJ/mol, calculated from the experimentally measured KD) (Figure 3B), suggesting a significantly increased affinity to human ACE2; the other mutants showed a similar ΔG compared to the prototype (Figure 3B). Compared to the KD (14.7 nM) of the prototype RBD, the equilibrium dissociation constant (KD) of V367F mutant was calculated as 0.11 nM (Figure 3C), which was two orders of magnitude lower than the prototype strain, indicating a remarkably increased affinity to the human ACE2 receptor. In comparison, the bat CoV RBD (strain RaTG13, with the highest genome similarity) showed a much lower binding affinity (KD=1.17mM; ΔG=−17.4kJ/mol) to human ACE2 than the pangolin CoV (KD=1.89μM; ΔG=−33.9kJ/mol). The affinity of the pangolin CoV to human ACE2 was lower than the SARS-CoV-2 prototype strain (KD=14.7nM; ΔG=−46.5kJ/mol) (Figure 3B, 3C).

**Fig. 3:**
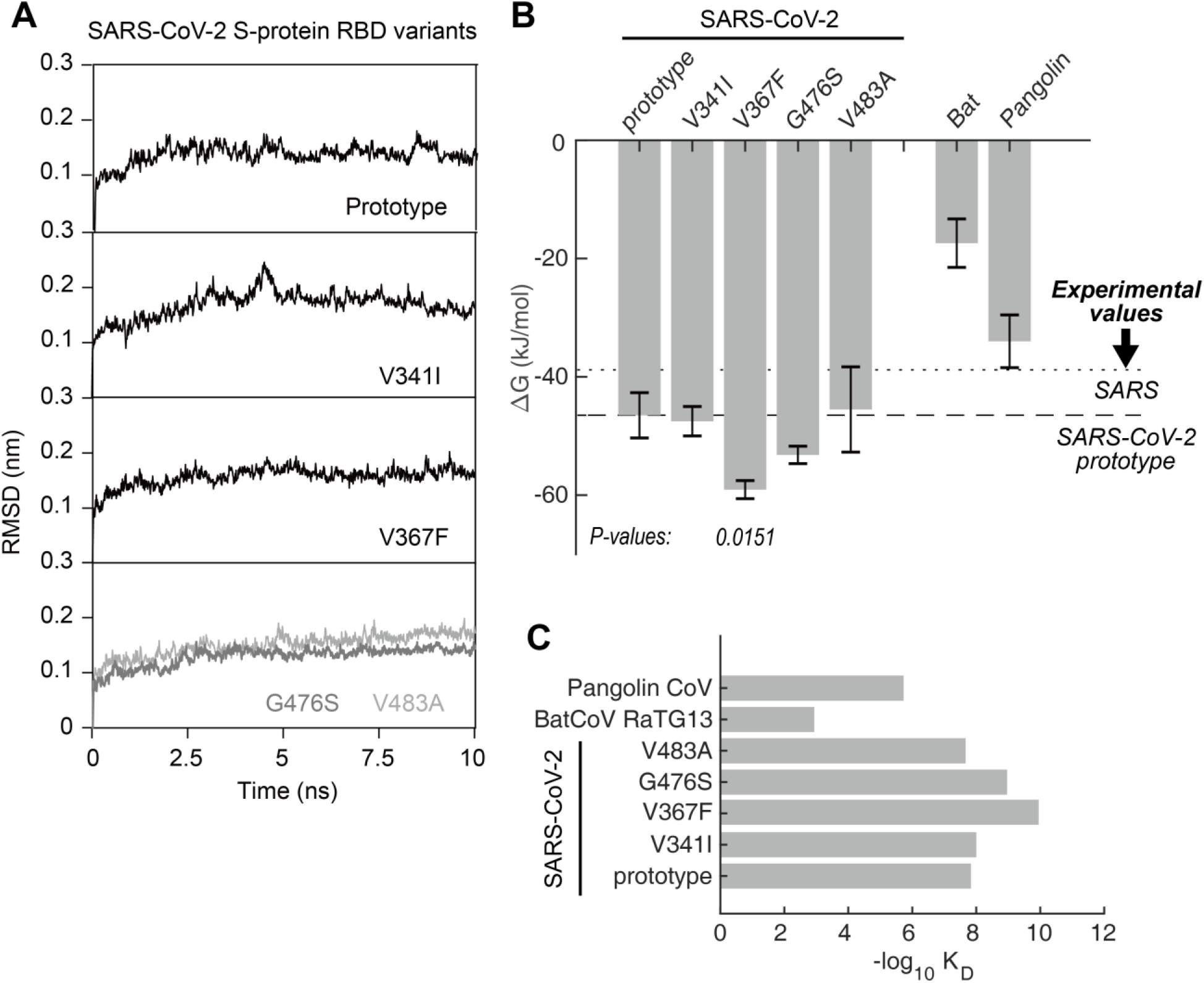
Binding free energy calculated for the SARS-CoV-2 S-RBD to human ACE2 receptor. **(A)** RMSD of typical MD trajectories of the SARS-CoV-2 prototype and the mutants. **(B)** Comparison of the binding free energy (ΔG) of the RBDs of human SARS-CoV-2, Bat CoV, and pangolin CoV, with the human ACE2. Note, the ΔG is inversely proportional to the binding affinity. Data are presented as mean ± SD. *P*-values were calculated using single-tailed student t-test. The *P*-values are shown for those with *P* < 0.05. The ΔG calculated from experimental K_D_ values of SARS and SARS-CoV-2 prototype are marked in dotted and dashed lines, respectively. **(C)** Comparison of the equilibrium dissociation constants (K_D_) as calculated with the ΔG.

### 2.4 Structural basis for the increased affinity of the SARS-CoV RBD mutants

To elucidate the structural basis of the increased affinity of V367F mutant, we investigated the dynamics of the residues in these structures in greater detail. The binding surface of the RBD to ACE2 is largely arrayed in random coil conformation, which lacks structural rigidity. Logically, a firm scaffold should be necessary to maintain this conformation of the interaction surface, and thus may facilitate the binding affinity. The beta-sheet structure scaffold, centered by residues 510-524 (Figure 4A, marked as red), apparently provides this rigidity. “Higher affinity” mutant V367F showed a considerable decrease of the RMSF (Root Mean Square of Fluctuation) at this region, demonstrating a more rigid structure; this was not observed for other mutants (Figure 4B). Coincidentally, the substitutions that accounts for the affinity increase of V367F are all located near this fragment. Indeed, residues 475-485, which is a random coil near the binding site, showed a remarkably higher RMSF for the “similar affinity” mutants, in contrast to the “higher affinity” V367F mutant (Figure 4B). Moreover, the “higher affinity” mutant V367F exhibited a general decreased ΔG in the binding site region in contrast to the “similar affinity” mutants (Figure 4C).

**Fig. 4:**
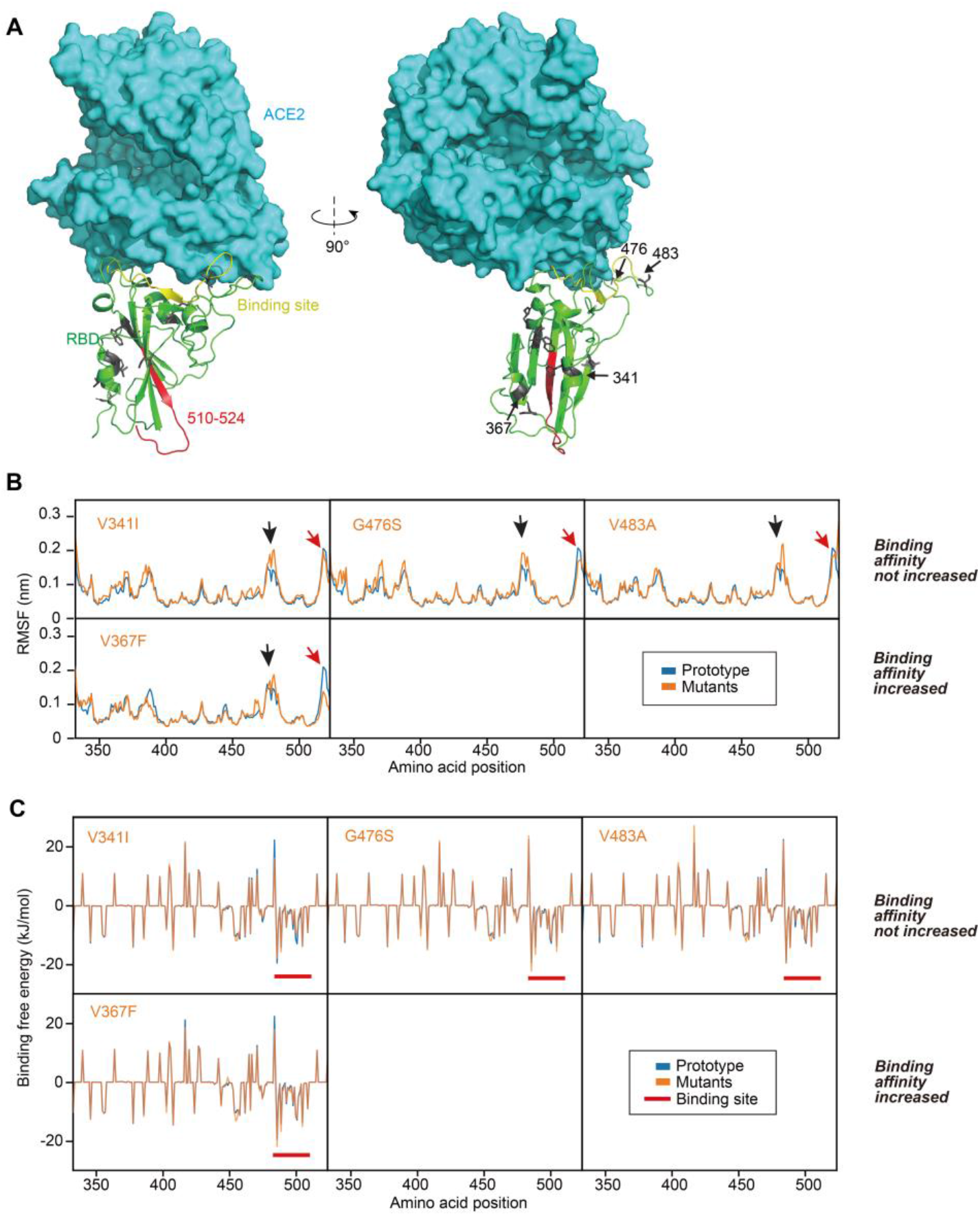
Structural analysis of RBD mutants and the effects on their binding affinity. **(A)** The binding surface and interaction of the RBD to ACE2, with the locations of the mutant amino acids noted. Beta-sheet structure scaffold was centered by residues 510-524 (in red). **(B)** Root Mean Square of Fluctuation (RMSF) of the mutants were compared to that of the prototype. Red arrows denote the fragment of residues 510-524. Black arrows denote the fragment of residues 475-485. **(C)** Contribution of each amino acid to the binding free energy. Red bars denote the binding site.

### 2.5 Experimental validation of the enhanced affinity and infectivity of the V367F mutant

As of March 31, 2020, among all of the mutants with dominant mutations in spike RBD (>10 isolates), most mutations resulted in a substitution of amino acids with similar properties. V367F is the only mutant with higher binding affinity, as calculated by MD simulation (Figure 3). Therefore, the binding affinity and the infectivity of V367F mutant were further validated experimentally. First, we performed experiments to assess the binding affinity in vitro with a receptor-ligand binding ELISA assay using purified S proteins and human ACE2 protein. The result showed that the V367F mutation significantly lowered the ED_50_ concentration (ED_50_ = 0.8±0.04 μg/ml), as compared to the prototype (ED_50_ = 1.7±0.14 μg/ml) (Figure 5A). This demonstrates that the V367F mutant has higher affinity with human ACE2 than the prototype. Second, we performed surface plasmon resonance (SPR) experiments, which yielded the same conclusion: the prototype had a KD of 5.08 nM, compared to the V367F mutant with a KD of 2.70 nM (Figure 5B). Additionally, we performed a virus-cell interaction experiment to investigate the invasion efficiency of S proteins using a HIV-backbone pseudovirus assay. Same amounts of S protein-containing pseudovirus were subjected to infection of ACE2-overexpressed Vero and Caco-2 cells. A higher infection efficiency is represented by the increased copy number of the integrated lentivirus genome. At 24 hours post-infection (h.p.i.), the V367F mutant pseudovirus showed 6.08x higher copy numbers than the prototype in Caco-2 cells (P<0.01). At 48 h.p.i, the V367F mutant pseudovirus showed 4.38x and 3.88x higher copy number than the prototype in Vero and Caco-2 cells, respectively (P<0.01) (Figure 5C). Therefore, the computation, protein and cell validations were consistent with each other: the V367F mutant had enhanced affinity and infectivity.

**Figure 5.**
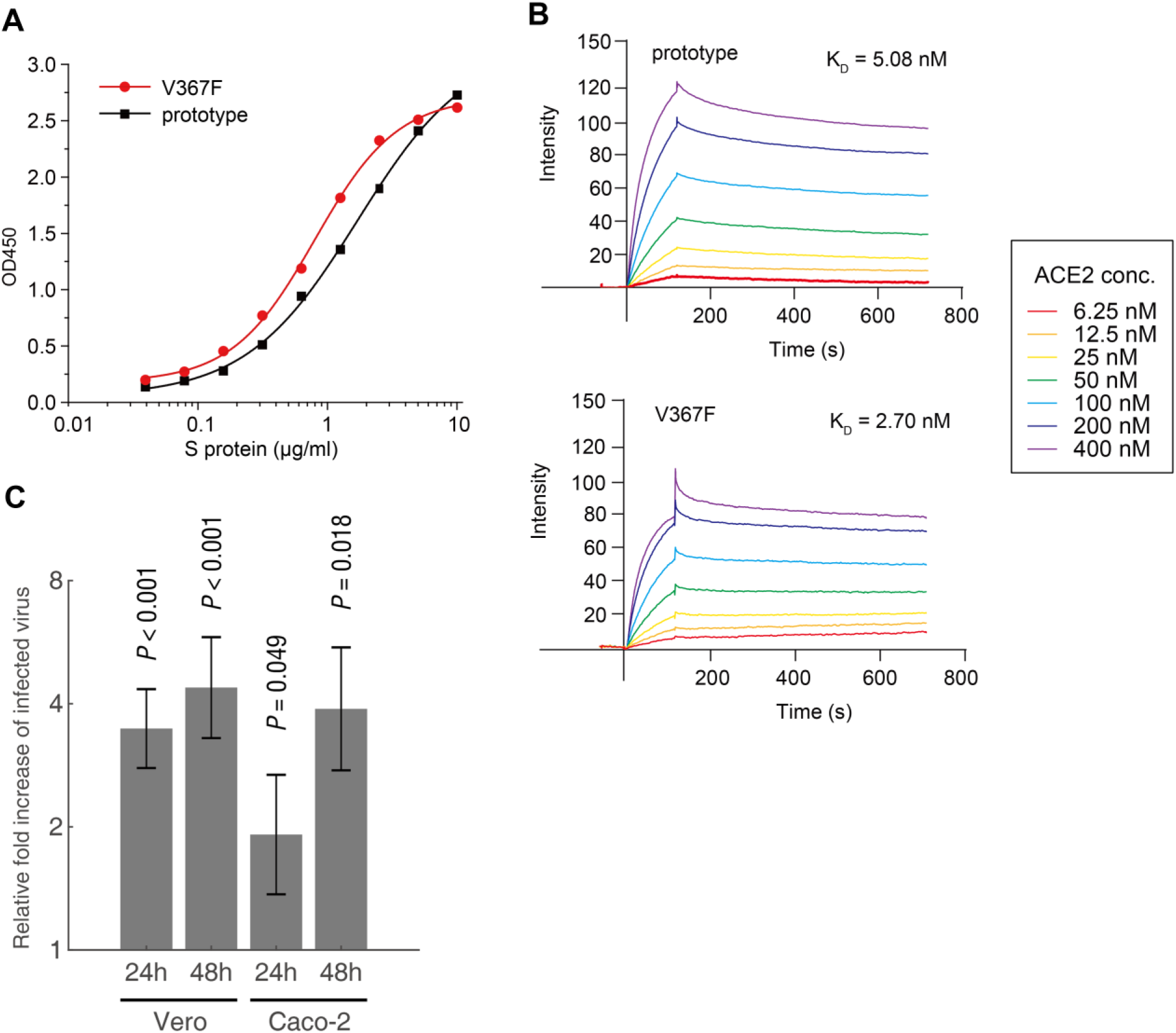
Experimental validation of the enhanced affinity and infectivity of the V367F mutant. **(A)** Comparison of the binding affinity of prototype S protein and V367F mutant to human ACE2 receptor by ligand-receptor binding ELISA assay. **(B)** Comparison of the binding affinity of prototype S protein and V367F mutant to human ACE2 protein by SPR. **(C)** Quantification of the genome copy number of the V367F mutant vs. the prototype using pseudovirus infection assay. The relative fold increases of viruses infecting the cells are shown by the pseudoviral DNA copy number of V367F mutant in both Vero and Caco-2 cells at 24h and 48h post-infection. Experiments were performed in triplicates, and the P-values were calculated using two-tailed t-test for two samples with different variances.

### 2.6 The convergence of SARS-CoV-2 RBD and D614G mutations in dominant mutation isolates

The D614G mutation is located in the S1 region and is outside the RBD of the SARS-CoV-2 spike protein. It has been confirmed that the D614G mutant increases virus infectivity by elevating its sensitivity to protease, which mutant has spread widely (23). Among all of the dominant RBD mutations (>10 isolates), V367F, A344S and G476S mutations were detected along with the D614 prototype during the early transmission phase. Later, the D614G mutant emerged along with these RBD mutants (Table 1).

The V367F mutants were initially discovered in January in Hong Kong. Afterwards, the V367F mutants emerged mainly in Europe, including United Kingdom, the Netherlands, Austria, and Iceland, as well as in the USA, Australia, and China. The D614G+V367F dual mutant was initially emergent in March in the Netherlands (Supplementary Table 1).

The phylogenetic analysis of the V367F mutant genomes during the early transmission phase showed that V367F mutants clustered more closely with the SARS-CoV-2 prototype strain in L clade. Intriguingly, all of the dual-mutation variants (V367F + D614G) emerging later formed a distinct sub-cluster in GH, GR, G clades, separate from the protype sub-cluster (**Figure 6)**(27). This indicates that the V367F mutation may have evolved along with D614G mutation, suggesting a synergistic effect of increased infectivity. Compared with all the single mutation (V367F) variants, dual mutation (V367F + D614G) variants were emergent in both GH and GR clades. Multiple dual mutation (V367F + D614G) variants located in different clades had mutations which were not detected in early V367F mutants. This indicates that the dual mutations may have evolved through several individual evolution events (**Figure 6)** (23, 28).

**Figure 6.**
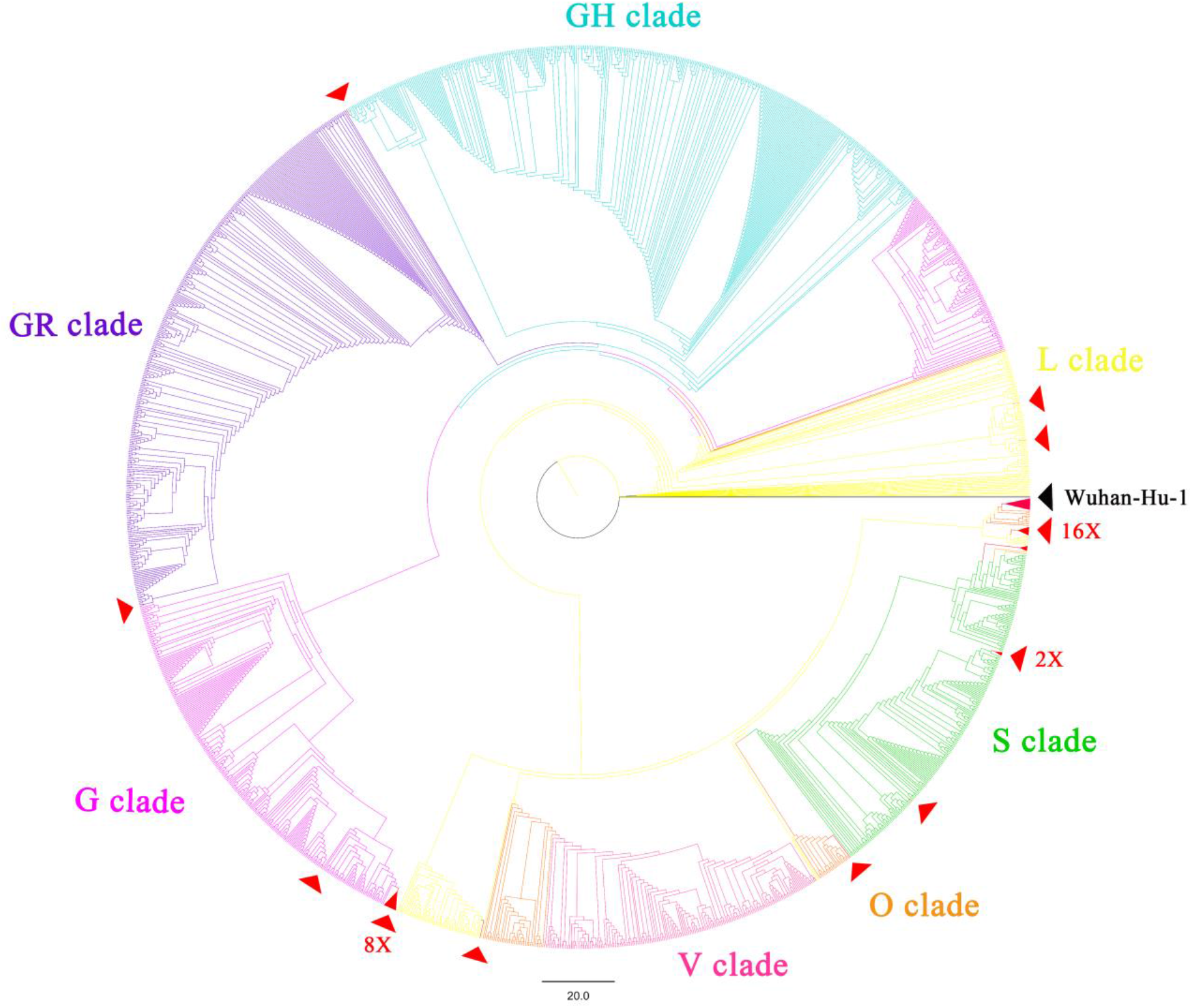
The whole genome phylogenetic analysis of the SARS-CoV-2 variants and mutants. (December 2019 through March 2020). The whole genome phylogenetic tree was constructed by IQ tree 2.02 using the maximum likelihood method with GTR+F+R3 model and 1000 bootstrap replicates, and applying default parameters. All 34 V367F mutants were included. For reference, the branch of Wuhan-Hu-01 is marked in black. V367F mutants are marked in red; sampled referenced sequences are annotated in different colors by clades using Figtree 1.4.4 (https://github.com/rambaut/figtree/releases).

## 3. Discussion

Due to the challenging and on-going pandemic and given the evolving nature of the SARS-CoV-2 virus globally, identifying changes in viral infectivity may prove crucial to containing the COVID-19 pandemic. Quarantine policies need to be adapted with respect to the viral infectivity change. It is always a dilemma of quarantine and economy. Any government would balance the lost due to the quarantine lockdown versus the lost due to the disease. Numerous models have been raised to estimate the two costs. For example, if the viral strain is more infectious, more stringent lockdown measure would be expected. This report provides computational and experimental insights into the functional outcome of mutations in RBD. As RBD mutations are under positive selection pressure, we identified the mutants that acquired increased binding affinity to human ACE2 receptor and presumably higher infectivity for human cells.

First, our analysis of molecular dynamics simulation indicated a remarkable enhancement of the affinity efficiency of multiple mutant S proteins. Compared to the prototype strain Wuhan-Hu-1, the ΔG of mutants decreased ~25%. These mutants bind to ACE2 more stably due to the enhancement of the base rigidity of the S protein structure. Potential and recent animal-to-human transmission events of SARS-CoV-2 may explain the strong positive selection and enhancement of the affinity during the pandemic. Ongoing adaptation to transmission and replication in humans, including mutation events in the RBD may boost the binding affinity and lead to an increase in the basic reproduction number (R0), and in theory, further enable human to human transmission.

The origin of this virus has been of considerable interest and speculation since the outbreak. Due to the high sequence similarity of the bat SARS-like CoV genome and the pangolin CoV RBD sequence to the SARS-CoV-2 sequences, these hosts were thought to have initiated the infection in humans(5, 6, 30) Our results provide more clues to support this hypothesis. Our results suggest that the binding energy of the bat SARS-like CoV RBD is too high to bind human ACE2 effectively (KD in millimolar range). In contrast, the pangolin CoV showed a binding KD for human ACE2 at the micromolar range, just ~6x higher than that of human SARS virus (K_D_ = 0.326μM)(Figure. 3), which indicates that the pangolin CoV has the potential to infect humans in unprotected close contact. Alignments of the genomic sequences of SARS-CoV-2 and pangolin CoV viruses suggest recombination events, particularly in the RBD domain, between pangolin and bat viruses (28).

The V367F mutants were found during the early transmission phase. As RBD is conserved in SARS-CoV-2, the coincidence of V367F mutants across large geographic distances indicates that this mutation is more robust and that these variants originated as a novel sub-lineage, given the close isolation dates (January 22 and 23, respectively). An alternate view is that asymptomatic individuals with the same mutation were “super-infecting” travelers. Along with the epidemiological data, mutation surveillance is of critical importance as it can reveal more exact transmission routes of the epidemic and provide early warnings of additional outbreaks. Emergence of SARS-CoV-2 mutants in Hong Kong, France, and other countries with RBD mutations allowing higher binding affinity to human ACE2 receptor suggests an increased risk of emergence of other mutants with “high-affinity” and increased infectivity during a sustained pandemic of COVID-19, particularly if effective precautions are not completely implemented. By performing assays comparing the prototype spike protein to the V367F mutation containing counterpart, we confirmed the significantly enhanced binding affinity and likely higher infectivity of the V367F mutant, and showed that the mutation stabilizes the RBD structure.

By tracking mutation types in the SARS-CoV-2 spike RBD, most of the dominant RBD mutants were first detected within the prototype D614 strain, and then together with the G614 variants. This is possibly due to multiple individual recombination events between the D614 mutants and G614 mutants, although it is difficult to determine the exact position and time point due to the high sequence identities among SARS-CoV-2 mutants. D614G is distinct from the RBD mutations – it is not located in the RBD but enhances viral infectivity by elevating its sensitivity to protease and increasing stability (12, 13, 23). Furthermore, D614G and V367F may function independently and have synergistic effects on viral infectivity. Recombination is known to play an important role in natural coronavirus evolution, which may contribute to the convergence of dual enhancing mutants (D614G + V367F). The phylogenetic and substitution analyses indicated that the divergent clade of dual mutation mutants were possibly from multiple individual recombination events which introduced new mutations. Whether these new mutations may produce more aggressive viruses remains further research. More attention should be paid to the risk of the advantage accumulation evolution through recombination among the variants. In this case the recombinants may be more infectious and also better at immune escape.

The S protein is also important for antigen recognition. By tracking dominant RBD mutants up to March 31, 2020, multiple mutations with more than 10 isolates putatively related to host receptor binding and affinity have been detected in this study. The equivalent positions in more than 500 SARS and MERS genomes were compared. Among them, V483A in the MERS-CoV and N439K in SARS-CoV resulted in weakly reduced host receptor binding and altered antigenicity, revealing possible immune escape driving virus evolution (32). Given the alanine shares the similar chemical and structural properties with serine, A344S variant is expected to have similar affinity to human ACE2 with the prototype strain. However, since the RBD contains important antigenic epitopes, RBD mutations, especially those that change the amino acid properties, may weaken the binding affinity of any antibody raised against the prototype strain. This may lead to decreased vaccine efficacy and should be further studied.

In summary, we have found 302 RBD mutants clustering into 96 dominant mutant types during the early transmission phase. Among the six dominant mutation types, V367F mutants that emerged in Asia and Europe displayed enhanced structural stability of the spike protein along with higher binding affinities to the human ACE2 receptor. V367F mutants have been continuously circulating along with the D614G mutants. This indicates that the V367F mutants are stable and may have acquired increased infectivity for humans during the COVID-19 pandemic. The emergence of dual mutants (V367F+D614G) possibly due to the recombination may be a hint of emergence of other variants with increased infectivity or with enhanced escape from the host immune response. These findings support the continuing surveillance of spike mutations to aid in the development of new COVID-19 drugs and vaccines.

## 4. Methods and materials

### 4.1 Genome sequence dataset in this study

Full-length gene sequences of SARS-CoV-2 were downloaded from the NCBI GenBank Database, China 2019 Novel Coronavirus Resource (https://bigd.big.ac.cn/ncov), and GISAID EpiFluTM Database (http://www.GISAID.org). 34,702 SARS-CoV-2 full-genome sequences isolated during early transmission phase (sample collected before March 31, 2020) were downloaded and the sequences with amino acid mutations in S protein and RBD region were parsed. The genome sequences with either the V367F mutation or the V367F/D614G dual mutations in the S protein RBD were screened and analyzed in this study (Supplementary Table 1). For evolution analysis, 1,753 full genome sequences from each Nextstrain (27) clade were filtered and cluster random sampled from GISAID EpiFluTM Database.

### 4.2 Mutation analyses and phylogenetic analyses

Alignment of S protein sequences from different sources and comparison of ACE2 proteins among different species were performed using MAFFT version 7, with default parameters (https://mafft.cbrc.jp/alignmeloadnt/server/) and BioEdit (33, 34). Selection pressure analyses were conducted using the Nei-Gojobori method(jukes-cantor model) with Mega X(version 10.0.2)(20, 21) . Substitution mutation analyses of mutants are performed compared with Wuhan-Hu-01 sequence using BioAider (version 1.314)(17). Phylogenetic analyses were conducted with Iqtree2, applying the maximum-likelihood method with 1000 bootstrap replicates using GTR+F+R3 model by MFP ModelFinder (35). The trees were annotated and modified using Figtree (version 1.4.4) (https://github.com/rambaut/figtree/releases).

### 4.3 Molecular dynamics (MD) simulation

The complex structure of the SARS-CoV-2 S-protein RBD domain and human ACE2 was obtained from the Nation Microbiology Data Center (ID: NMDCS0000001) (PDB ID: 6LZG) (https://www.rcsb.org/structure/6LZG)(25). Mutated amino acids of the SARS-CoV-2 RBD mutants were directly replaced in the model, and the bat/pangolin CoV RBD domain was modelled using SWISS-MODEL(26). Molecular dynamics simulation was performed using GROMACS 2019 with the following options and parameters: explicit solvent model, system temperature 37°C, OPLS/AA all-atoms force field, and LINCS restraints. With 2fs steps, each simulation was performed at 10ns, and each model was simulated three times to generate three independent trajectory replications. Binding free energy (ΔG) was calculated using MM-PBSA method (GitHub; https://github.com/Jerkwin/gmxtool), with the trajectories after structural equilibrium assessed using RMSD (Root Mean Square Deviation)(24,36). To calculate the equilibrium dissociation constant (KD) and ΔG, the formula ΔG=RTlnK_D was used. Estimated ΔGs of the RBD mutants were normalized using the ΔG of the prototype strain which was derived from experimental data.

### 4.4 Expression of recombinant S protein mutant

The SARS-CoV-2 prototype S gene was cloned into pNPM5 vector (Novoprotein, NJ, USA), and fused with a C-terminal His6-tag. The V367F mutation was introduced using site-directed mutagenesis, consistent with the nucleotide sequence of the actual isolate. These two constructs were transfected into HEK293 cells using polyethyleneimine. Since the S protein included a signal peptide in its N-terminal 14 amino acids, it was secreted into the medium. The expressed proteins were purified from filtered cell supernatants with a Ni-NTA column. Eluted protein solution was then dialyzed against PBS (pH7.4) for subsequent assays.

### 4.5 Ligand-receptor binding ELISA assay

Human ACE2 protein was immobilized onto a microtiter plate at 5 μg/ml (100μl/well). Each S protein (prototype and V367F) was added as a ligand at different concentrations, ranging from 0.03 μg/ml to 10 μg/ml, and then incubated for 2 hours at 37°C to allow receptor-ligand interaction. The ligand / receptor mixture was then washed three times. 100μl of HRP anti-His Tag Antibody (BioLegend, USA) (diluted 1:20000) was added to each well, and allowed to react for 1 hour. After three washes, the signal was visualized using TMB solution (Sigma-Aldrich, USA) and a microtiter plate reader recording OD450.

### 4.6 Surface Plasmon Resonance (SPR) experiments

The SPR experiments were performed in a BIAcore T200 instrument (GE, USA). For this, the SARS-CoV-2 S-proteins, either prototype or V367F, were immobilized on the Sensor Chip NTA (GE, USA), according to the manufacturer’s protocol. Human ACE2 protein was injected in each experiment in seven concentrations (6.25, 12.5, 25, 50, 100, 200, and 400 nM). For each cycle, the absorption phase lasted for 120 seconds and the dissociation phase lasted for 600 seconds. After each cycle, the chip was regenerated using 350mM EDTA and 50mM NaOH wash for 120 seconds, so that the chip was ready for the next round of S-protein immobilization. Blank controls with 0 nM ACE2 were performed, with the blank signals subtracted from the cycle signals. All experiments were performed at 37°C. KD values were calculated by fitting the curves using the software provided with the instrument.

### 4.7 Production and titration of SARS-CoV-2 pseudoviruses bearing V367F S protein

The full-length S gene of SARS-CoV-2 strain Wuhan-Hu-1(NC_045512.2) was cloned into a SARS-CoV-2 Spike vector (PackGene, Guangzhou, China), and confirmed by DNA sequencing. Plasmid SARS-CoV-2 Spike (G1099T), incorporating the V367F mutation in the S gene, was constructed by site-directed mutagenesis using the ClonExpress MultiS One Step Cloning Kit (Vazyme), as per the manufacturer’s protocol.

Generation of SARS-CoV-2 S HIV-backbone pseudovirus was done as previously described with some modifications(36, 37). Briefly, 293T cells, at about 70-90% confluence, were co-transfected with 9 ug of the transfer vector (pLv-CMV), 6 ug packaging plasmid (psPAX-lentiviral), and 6 ug envelope plasmid (pCB-spike or pCB-spikeV367F). Pseudoviruses were harvested at 48 h post-transfection, using a 2.5 ml sterile syringe and subsequently filtered into either Falcon or microcentrifuge tubes via a syringe driven 0.45 μm filter. Virus titration was measured by RT-qPCR targeting the WPRE gene of pseudoviruses and using the Hifair® Ⅲ One Step RT-qPCR SYBR Green Kit (Yeasen), as per the manufacturer’s protocol. After reverse transcription (10 min at 42°C) and initial denaturation (5 min at 95°C), PCR amplification was performed for 40 cycles (15 s at 95°C, 60 s at 60°C). Primers were as follows: WPRE-F, 5’- CACCACCTGTCAGCTCCTTT-3’; WPRE -R, 5’- ACGGAATTGTCAGTGCCCAA-3’.

### 4.8 SARS-CoV-2 spike-mediated pseudovirus entry assay

To detect S variants mediated viral entry, VERO E6 and Caco2 cells (5×10 ^4^) grown in 24-well plates were respectively infected with either 20 MOI of S-V367 or S-F367-bearing pseudovirus (1×10^7^ pseudovirus copies per ml). The cell medium was replaced by 500 μl fresh DMEM medium 6h post-infection. Relative fold increase of infected virus titers was calculated according to the WPRE DNA copy number of the lentiviral proviruses measured by TB Green® Premix Ex Taq ™ II (Takara) at 24 and 48 h post-infection. Data from all of the samples were obtained from three independent experiments, and each sample was tested in triplicate.

## Acknowledgments

We gratefully acknowledge the authors, originating and submitting laboratories of the sequences from GISAID’s EpiFlu™ Database on which this research is based. All submitters of data may be contacted directly via **www.gisaid.org.**

Data and acknowledgement of GISAID sequences are available in Supplementary Table 2.

## Supplementary data

**Supplementary Table 1:** Meta data of the isolates with V367F mutations in the RBD of spike glycoprotein(January through March, 2020)

**Supplementary Table 2:** Acknowledgement table for GISAID sequences

**Supplementary Figure 1:** Multiple alignments of the RBD amino acid sequences. SARS-CoV-2 Wuhan-Hu-1, the first reported genome, is used as reference. Bat and pangolin SARS-like coronaviruses are also included. Amino acid substitutions are marked. Dots indicate identical amino acids.

